# A male-transmitted B chromosome undergoes strong meiotic drag in females

**DOI:** 10.1101/2025.05.20.655026

**Authors:** J. Cummings, E. Garman, J. Solomon, K. Soriano Martinez, P. M. Ferree

## Abstract

Many organisms carry extra, non-essential chromosomes known as B chromosomes (Bs), which are selfishly transmitted at super-Mendelian levels to offspring. This heightened transmission, termed drive, occurs during gametogenesis, usually in one of the two parents. In some cases, Bs can experience an opposing process, drag, which reduces their transmission. If these processes occur together in the same organism, one in each parental sex, then they may facilitate the spread of Bs while countering their accumulation in the genome to harmful levels. While previous studies have elucidated mechanistic aspects of B drive, little is known about drag or other factors that govern the inheritance of these selfish genetic elements. Here we examined the inheritance of Paternal Sex Ratio (PSR), a single-copy B in the jewel wasp, *Nasonia vitripennis*, which is transmitted paternally to offspring. PSR drives by converting female-destined embryos into PSR-transmitting males. Using genetic manipulation, we produced exceptional PSR-carrying females, which were used to assess B transmission potential. We found that females transmit PSR at unexpectedly low levels compared to univalent chromosomes in other organisms. This reduced transmission stems from remarkable loss of PSR from the egg’s nucleus upon entry into meiosis, an effect that may be caused by an absence of microtubule-based spindle fibers in meiosis I-arrested wasp eggs. We also found that PSR is strictly limited to a single copy per genome, suggesting that two PSR copies are lethal during development. Our findings reveal the successful inheritance of this selfish B chromosome involves a restriction to a single copy and hidden female meiotic drag in addition to its strong paternal drive.

## INTRODUCTION

Mendelian inheritance predicts the fair transmission of differing alleles from a heterozygous parent to progeny. This fundamental pattern of inheritance relies on equal segregation of homologous chromosomes into sperm or egg during gametogenesis. However, thousands of plants, animals, and fungi carry non-essential chromosomes known as B chromosomes (Bs), many of which exhibit deviant transmission characteristics that are self-serving and, thus, evolutionarily advantageous for these selfish genetic elements [1][2][3]. The number of Bs typically ranges from 2 to 6 copies per species but some can reach as high as 30-50 copies [3,4], and within a given species, B copy number can vary greatly among individuals [5]. These peculiar dynamics and, more broadly, the long-term persistence of B chromosomes, are strongly influenced by two opposing phenomena known as *drive* and *drag* [6] While Bs normally are transmitted to progeny by both parents, they often exhibit super-Mendelian (*i.e.*, positively biased) transmission from one or the other parental sex. In many organisms, this biased transmission, known as drive, occurs during female meiosis and is influenced by certain properties of meiotic cell division. For example, in certain organisms such as the mottled grasshopper and the slim stem lily, Bs segregate more frequently into the egg-destined meiotic product because they tend to aggregate on the longer side of an asymmetrically shaped spindle apparatus during the first meiotic division [7,8]. Bs in the fruit fly *Drosophila melanogaster* also undergo drive during female meiosis [9], but presumably through a different mechanism because the first meiotic spindle in this organism is not spatially asymmetrical [10][11]. Regardless, these Bs are preferentially transmitted, resulting in progeny possessing as many as 12 B copies per individual [9]. In contrast, in this same insect, Bs are transmitted by the male parent at lower-than-Mendelian levels [9] through a process referred to as drag [12]. In such cases where drive and drag occur in the same organism, one in each parental sex, these processes may help to maintain B copy number at levels that prevent B loss and simultaneously prevent B copy number from reaching levels that are deleterious to the organism. It is universally likely that specific aspects of gametogenesis that differ between the sexes lead to drive in females and drag in males in species like *D. melanogaster*, or vice versa in other organisms, and the interplay of drive, drag, and perhaps other unknown factors governs the overall success of B inheritance. While recent studies have helped elucidate mechanistic aspects of B drive in a number of organisms including *D. melanogaster* [9], rye [13] and maize [14], very little is known about the cellular features that underlie drag or otherwise delimit B transmission.

The jewel wasp, *Nasonia vitripennis*, carries a selfish B chromosome known as Paternal Sex Ratio (PSR), which exhibits exceptional but incompletely understood transmission characteristics. In wild caught isolates and laboratory stocks, PSR is found only in the male sex, and it is invariantly present in single copy [15]. Because PSR is only found in males, it is transmitted exclusively by father to offspring via the sperm. PSR’s drive involves a trans-acting effect that causes elimination of the sperm’s standard genome, but not PSR itself, during the first mitotic division following fertilization [16,17]. The reproductive mode of *N. vitripennis* and all other hymenopteran insects (including all wasps, bees and ants) is haplo-diploidy, whereby males are haploid and develop from unfertilized eggs having a single set of chromosomes, and females are diploid, arising from fertilized eggs having two chromosome sets. By eliminating the sperm’s hereditary material, female-destined embryos are transformed into PSR-carrying (PSR+), and thus -transmitting, males. Based on previous genetic evidence, this paternal genome elimination (PGE) event and the resulting female-to-male conversion are highly effective, occurring in over 99% of eggs fertilized by PSR+ sperm [18]. Given the strength of this effect, and because the fertilization rate in this organism is 80-90%, the result is the production of all-male broods, with the great majority of males being PSR+. Due to the remarkable effect of PSR on the wasp’s reproduction and sex ratio, this B has been referred to as the most extreme selfish genetic element currently known [19].

The exceptional inheritance characteristics of PSR beg several important questions. First, why is the transmission of this B restricted to the male sex? Part of the answer is likely that PSR’s extremely strong female-to-male conversion effect leads to this B invariantly being in the male sex. It is formally possible that PSR could be transmitted maternally, or even undergo meiotic drive like other Bs, if it were to find itself in females. Alternatively, there may be unknown fitness costs to females that carry PSR, or intrinsic cellular barriers to PSR transmission during female gametogenesis. It is noteworthy that unlike haploid male wasps, which produce sperm entirely through mitotic division [20], egg production in diploid females involves standard meiosis (ref). This difference between the sexes opens the possibility that PSR may encounter different cellular influences when transmitted by each parent. A second important question is: why is PSR present only as a single copy in the wasp genome? This characteristic may result from a lack of opportunity for PSR to reach a higher copy number because it is not transmitted by both parents. Instead, there may be a genetic constraint to the B being present in two or more copies per genome.

We tested these ideas by experimentation with artificially produced, PSR+ females. Previously it was shown that expression of a PSR gene named *haploidizer* is required for PSR’s genome elimination activity [21]. When transcripts of this gene are targeted for degradation by systemic RNAi (sRNAi), PSR’s genome eliminating activity is suppressed, leading to the production of daughters that carry PSR [21]. Using these individuals, we found that PSR does not affect female fitness, and females are capable of transmitting PSR to F1 progeny. However, the level of maternal transmission is much lower than predicted for a univalent chromosome, compared to univalent segregation in *D. melanogaster*. This transmission reduction stems from PSR being lost from the maternal nucleus upon entry into the first meiotic division, before chromosome segregation. Thus, PSR experiences meiotic drag in females, which is a rare occurrence due to PSR’s strong female-to-male conversion activity that enables its transmission through males. Finally, crosses between PSR+ males and females produced numerous adult offspring with a single PSR copy but none with two PSR copies, suggesting that two-copy individuals die during development. Together our findings argue that the successful inheritance of PSR depends largely on the balance between its strong paternal drive and restriction to one copy, although the observed maternal drag, normally masked by strong drive, may help eliminate non-driving PSR variants and ultimately enhance wasp fitness.

## RESULTS

### PSR undergoes normal mitotic segregation in females and does not affect their fitness or fecundity

Previously, it was determined that genetically produced PSR+ females do not transmit PSR to their offspring [21]. To revisit this finding, we generated PSR+ females using the same approach (see Methods). After confirming the PSR-carrying status of these individuals with PCR (Figure 1A), we microscopically examined their ovaries. Like wild type ovaries, PSR+ ovaries contained developing oocytes that were morphologically normal (Figure 1B). Using DNA fluorescence in situ hybridization (FISH), we visualized PSR with a probe that is cognate to a high-copy satellite sequence uniquely located on PSR; this probe labels both arms of the B chromosome [22]. PSR was present in the nuclei of nurse cells in early-, mid- and late-stage egg chambers (Figure 1B). In these nuclei, PSR was amplified to high copy number, as indicated by the large region of FISH signal, like the wasp’s rDNA locus (Figure 1B). This pattern reflects the polyploidization of DNA in these cells [23]. PSR also appeared in the nuclei of the somatic follicle cells that surround each egg chamber, and as a small, single focus inside the oocyte’s germinal vesicle (Figure 1B). We saw no nuclei in the ovary that were devoid of PSR, indicating that PSR segregates effectively during the numerous mitotic divisions giving rise to the female’s germline and somatic cells. Additionally, PSR+ females were like wild type females in body size, longevity, egg production, and number of offspring produced per female (Figure 1C). Thus, PSR has no measurable effect on female fitness, and PSR+ females do not seem to suffer from potential aneuploidy or other chromosome abnormalities that may arise from the imperfect PGE suppression that was previously observed to occur in some young embryos because of the sRNAi treatment used to produce them [21].

**Figure 1.**
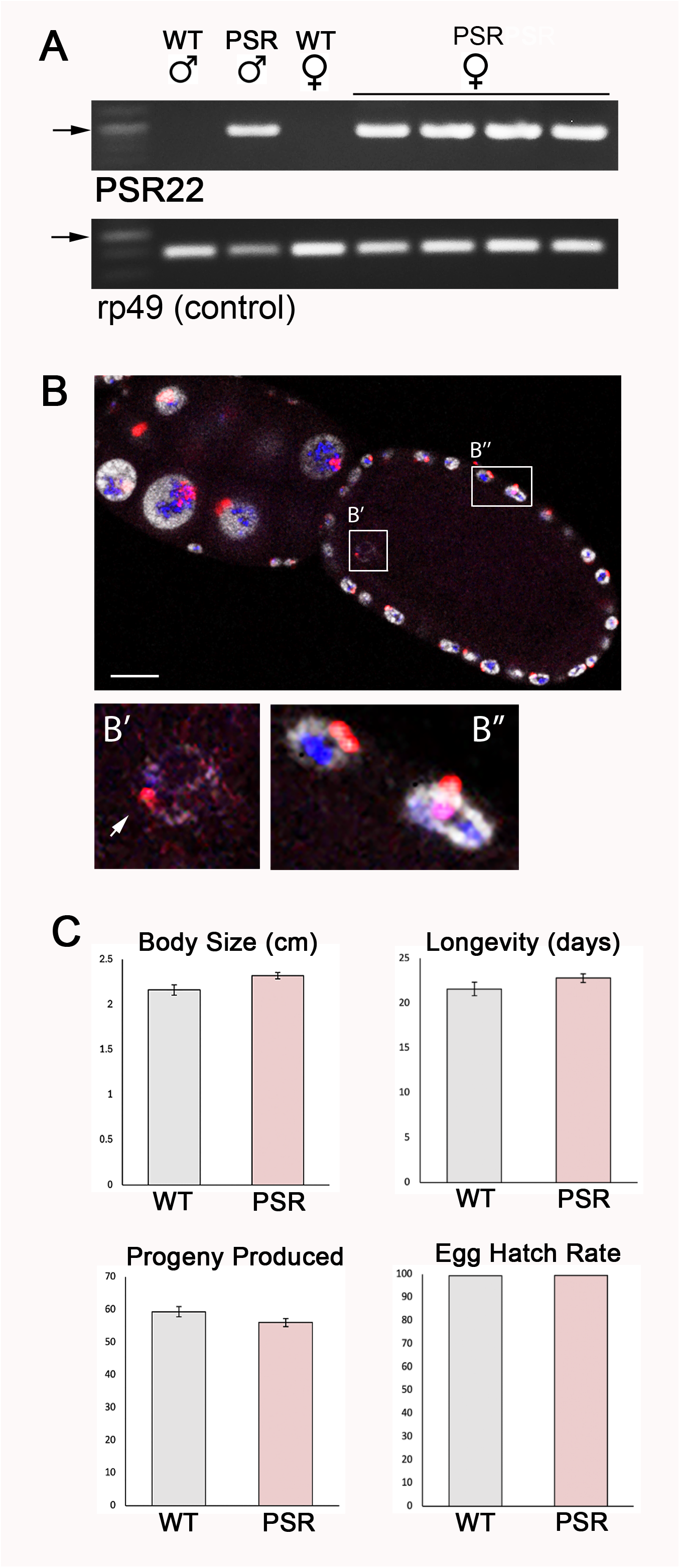
Genetically produced PSR+ females are similarly fit to wildtype females. (**A**) An agarose genotyping gel showing amplification of a ∼1 kilobase pair PSR-specific PCR product from the genomic DNA of F1 females produced by PSR+ fathers that were RNAi-treated for degradation of *haploidizer* transcripts. The arrows indicate a 1,000nt band in the size standard ladder. (**B**) A mid-stage ovary taken from a PSR+ female, when hybridized with a PSR probe cognate to the *haploidizer* gene, reveals the presence of PSR (red) in the polyploid nurse cells, the germinal vesicle (**B’**) and somatic follicle cells (**B’’**). rDNA is shown in blue, and DNA is grey. Scale bar equals 15 mM. (**B’**) and (B’’) are shown in higher magnification. The white arrow in (**B’**) indicates the single copy of PSR in the germinal vesicle’s chromatin. (**C**) Four different characteristics of fitness – body size, longevity, progeny produced per PSR+ female, and hatch rate for eggs laid by PSR+ females – are shown in comparison with the same characteristics for wildtype females. Standard error bars are shown for the first three characteristics.

### Females transmit PSR to progeny at lower-than-expected levels for a univalent chromosome

To assess the maternal transmission level of PSR, we used PCR to genotype F1 offspring produced by unmated, PSR+ females. Previous genetic and microscopic studies of univalent segregation in *D. melanogaster* were considered to estimate an expected level of PSR transmission to offspring from females. In the fruit fly, univalent chromosomes derived from fusions of two X chromosomes, two fourth chromosomes, or an arm from each of chromosomes 2 and 3 exhibited no substantial segregation defects during meiosis and were transmitted effectively to an estimated half of all eggs, when accounting for lethal progeny that do not receive a homologous counterpart [24]. In contrast to these univalent chromosomes in the fruit fly, PSR was detected in only 22 percent of offspring laid by unmated female wasps (Figure 2A). This transmission level was similar when PSR+ females were crossed with wild type males (Figure 2A), showing that fertilization does not affect the level of PSR transmission by females. These findings argue that, even though PSR segregates properly during mitosis in females and is present in the germinal vesicle of developing oocytes, this B is transmitted at a much lower level than previously observed for univalent chromosomes in the fruit fly. Notably, PSR+ females when crossed with PSR+ males produced all-male broods, like wild type females crossed with PSR+ males (Figure 2B). This finding demonstrates that the presence of PSR in the egg does not impact the PGE-inducing activity of the paternally transmitted PSR copy.

**Figure 2.**
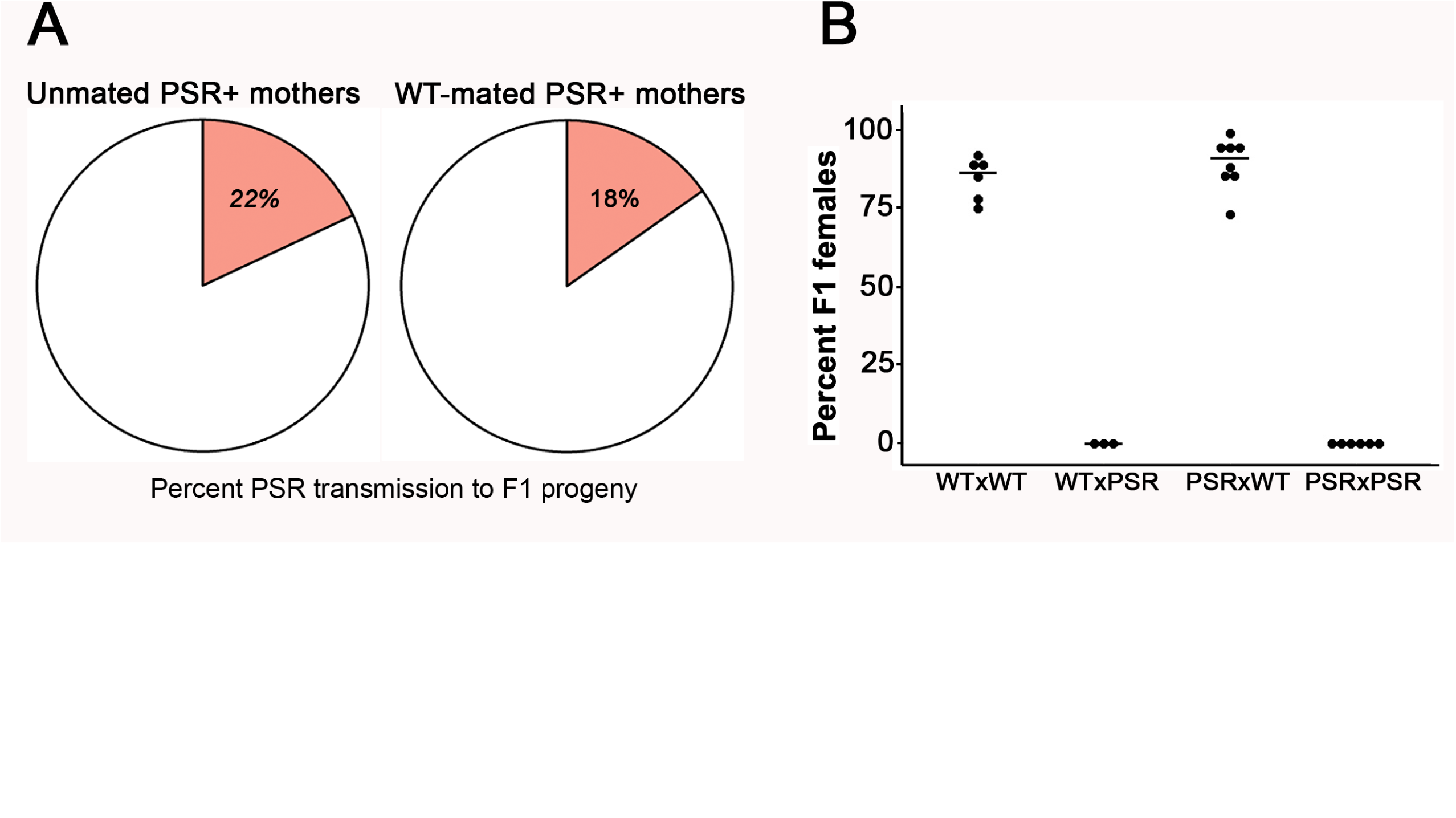
Mating status does not influence the low maternal transmission of PSR. (**A**) The percentage of F1 progeny inheriting PSR from unmated PSR+ mothers (left pie chart) or PSR+ mothers mated with wildtype males (right chart). The transmission levels for these conditions are very similar. (**B**) Maternally transmitted PSR does not cause PGE and sex ratio distortion, nor does it affect PGE caused by paternally transmitted PSR. Sex ratio values for adult F1 broods produced by PSR+ mothers are shown on the y-axis. The female x male genotypes are shown for each cross on the x-axis.

### PSR becomes lost from the nucleus upon entry into meiosis

To gain insight into the low maternal transmission of PSR, we microscopically inspected 1-2hr unfertilized embryos laid by unmated PSR+ females. By this time, the meiotic divisions within the egg’s cytoplasm have completed, and one of the four meiotic products which becomes the haploid embryonic nucleus has initiated syncytial mitotic divisions (Figure 3A; Figure S1). The other three meiotic products have become hyper-condensed polar bodies that cluster at a position near the inner side of the plasma membrane and do not undergo division (Figure 3A: Figure S1). In 11 percent of these embryos (n=15/133), the two sister PSR chromatids occupied either two polar bodies or one polar body and the mitotic cleavage nuclei, reflecting normal segregation patterns (Figure 3B). However, in the remaining 89 percent of embryos (n=118/133), either one or both sister PSR chromatids were found outside the mitotic nuclei or polar bodies, sometimes at a great distance from them in the cytoplasm (Figure 3B).

**Figure 3.**
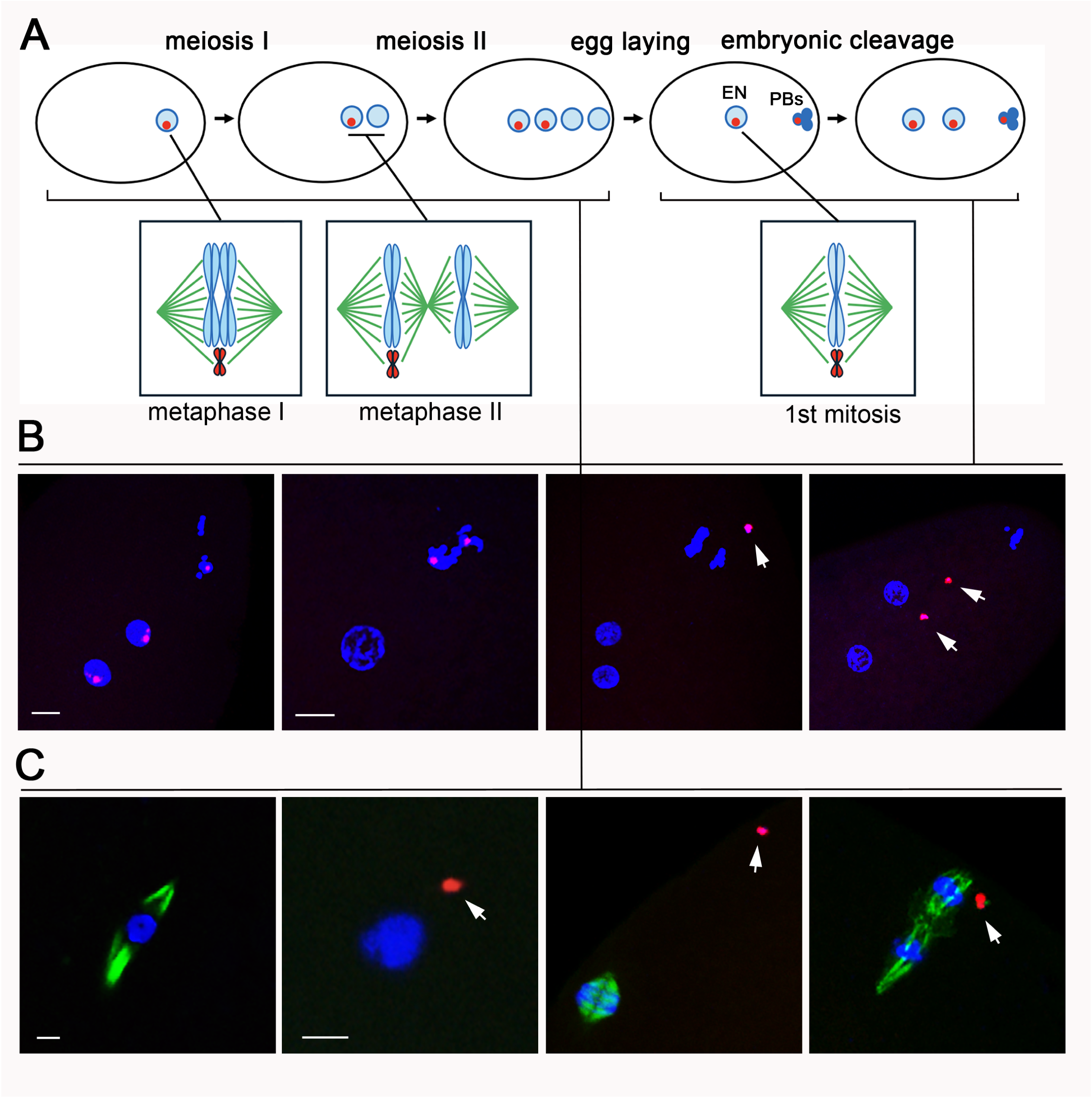
PSR is lost from the egg’s nuclear material upon entry into meiosis. (**A**) Schematization depicting PSR segregation during meiosis and the first embryonic mitotic division. Under this model, PSR is depicted to segregate normally to two of four meiotic products if there is a functional distributive system that ensures proper segregation of achiasmatic and univalent chromosomes. In the scenario shown, the sister PSR chromatid pair moves to the inner product at the end of meiosis I, leading to PSR being present in the egg’s nucleus and one polar body. An alternate scenario showing the sister PSR chromatid pair moving to the other meiotic product during meiosis I is shown in a supplemental figure (**Fig S1**). (**B**) The first embryonic mitotic division is shown in the panels, with different outcomes of PSR segregation. From left to right: PSR segregates to the two mitotic daughters and one polar body (*i.e.*, the scenario in the schematic); PSR (red) segregates to two of the three polar bodies (*i.e*., the scenario in Fig S1); loss of the sister PSR chromatid pair from the polar bodies; and loss of the PSR chromatids separately from the nuclei. Scale bars in the left panels equal 10 mM. White arrows indicate PSR sister pair or individual chromatids. (**C**) Panels show meiotic nuclei. Left-most panel shows the egg nuclear material arrested in meiosis 1. A bipolar spindle apparatus (green) is clearly visible. The next three panels are different stages of meiosis in *N. vitripennis*. From left to right: Entry into meiosis in a mature egg, where the chromatin is condensed but spindle fibers are not visible. PSR has already become lost from the meiotic nucleus; Metaphase of meiosis I, in which spindle fibers (green) are apparent and PSR has drifted from the spindle; Metaphase of meiosis II, showing a tripolar spindle and PSR at an adjacent position. White arrows indicate PSR. Scale bars equal 8 mM.

To identify when PSR becomes lost from the nuclei, we examined eggs undergoing meiosis. For comparison, the hereditary material of mature, unactivated *D. melanogaster* eggs is arrested at metaphase of meiosis I [25]. Initially, we examined mature fly eggs to confirm that our experimental conditions were conducive to visualizing the spindle apparatus. In these cells, a well-formed bipolar spindle could be clearly seen to emanate from the body of condensed meiotic chromosomes under fluorescent microscopy using an antibody against a-Tubulin (Figure 3C). In contrast, in mature, unactivated *N. vitripennis* eggs, the hereditary material is condensed but the spindle is not yet visible (Figure 3C; N=7/7). By this time, PSR has become detached from the nucleus (Figure 3C). Given our prior observation that PSR is present within the chromatin of the germinal vesicle in immature eggs before the initiation of meiosis, it stands to reason that PSR becomes lost from the nucleus upon entry into meiosis I. The displaced PSR chromatids were also observed at metaphase of the first and second meiotic divisions, when the spindle apparatus has clearly formed (Figure 3C). In contrast, the wasp’s essential chromosomes remained within the nuclei during all steps of meiosis and the early embryonic mitotic divisions (Figure 3C). Thus, the chromosomal loss is specific to PSR.

### PSR copy number is constrained to one per genome

Given that PSR is invariably present as a single copy per wasp genome in natural populations and laboratory stocks, we tested if wasps could carry two PSR copies. To do this, we crossed the genetically produced PSR+ females with PSR+ males and microscopically examined B chromosome copy number in their F1 progeny. Considering the measured maternal transmission rate of 22 percent, the fact that effectively all sperm of PSR+ males carry PSR [26], and ≥80% of eggs are fertilized [27], we estimated that 17.6 percent of F1 progeny from these crosses should carry two copies of PSR. We found that 16.7 percent of young F1 embryos (n=13/78) contained two clearly distinct and spatially separated PSR foci, one from each gamete, in each mitotic nucleus (Figure 4A, B). The remainder of F1 embryos (n=65/78) contained a single PSR focus in each nucleus, likely reflecting the copy delivered by the sperm (Figure 4A, B). We then assessed PSR copy number in F1 adults by examining the nuclei of their individualized, mature sperm cells. In all examined sperm from numerous males generated by different cross replicates (n=171/171 males from 7 different PSR+ x PSR+ crosses), we unambiguously observed only a single PSR copy (Figure 4C). Based on these results, it is likely that the somatic genotypes of the males that generated the sperm also carry a single PSR copy. The lack of any adults carrying two PSR copies suggests that this genotype may be lethal after early embryogenesis. Thus, there appears to be a strict limitation of PSR’s copy number to one.

**Figure 4.**
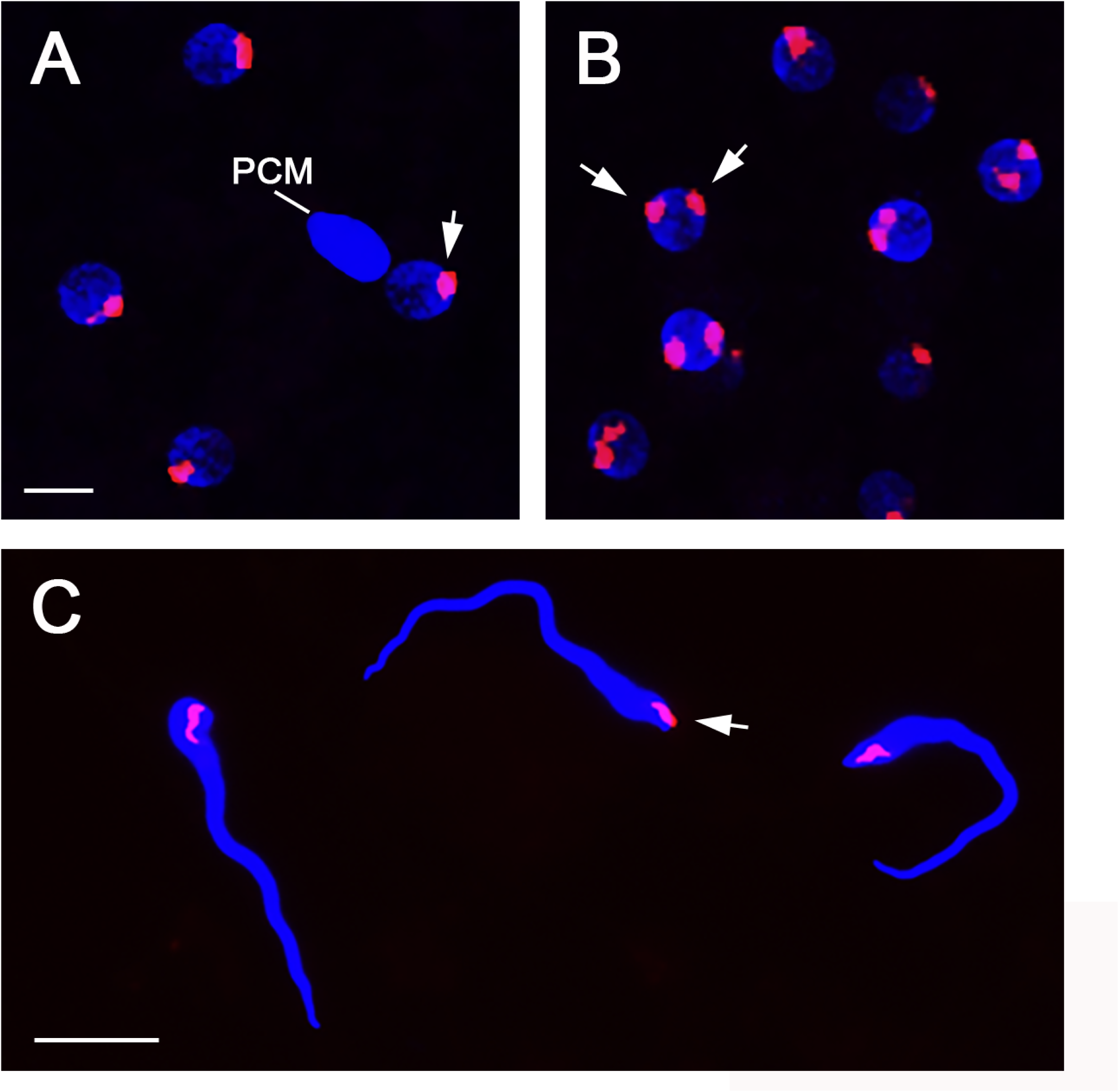
PSR is restricted to single-copy status in adult F1 males produced by PSR+ parents. (A) Nuclei in a 1-2hr old embryo produced by PSR+ parents. Each nucleus contains a single copy of PSR. (B) Nuclei in a slightly older embryo produced by the same parents as (A). Each nucleus contains two well-separated PSR copies (white arrows). (C) Nuclei of mature sperm cells from F1 adults produced by the same parents; each contains a single PSR copy (white arrow). Scale bars in (A) and (C) are 10 mM and 5 mM, respectively.

## DISCUSSION

In this study, we investigated two unexplored aspects of PSR inheritance by utilizing genetically produced PSR+ females. The first aspect is whether female wasps can transmit PSR. Normally this B chromosome is carried only by males due to its strong female-to-male-converting activity through PGE. Thus, the female sex is bypassed under normal circumstances. Just as there are no observable fitness differences between PSR+ and wild type males [28], the same is true for females. A previous study observed a lack of PSR transmission by females [21], which may have arisen from certain PCR conditions that were used for genotyping. Here, we unambiguously demonstrated with PCR and microscopic analysis that PSR+ females are indeed capable of transmitting PSR to progeny. However, PSR transmission by females is substantially lower than univalent chromosome segregation in *D. melanogaster*, which is Mendelian [24]. This reduced transmission is caused by the loss of PSR from the egg’s nuclear material as it enters meiosis. The magnitude of this loss measured microscopically matches the proportion of adult F1 progeny that did not receive a copy of PSR. We emphasize that PSR’s nuclear loss stands in contrast to the previously observed behavior of univalent chromosomes in *D. melanogaster*, which remain within meiotic nuclei and segregate properly to progeny [24]. Thus, our work reveals strong PSR drag in the female wasp, which we interpret is caused by an incompatibility of the B chromosome with some aspect of female meiosis in this organism.

What might underlie this incompatibility? In *D. melanogaster*, there exists a secondary mode of chromosomal segregation known as the distributive system, which is responsible for ensuring proper segregation of homologous chromosomes that do not undergo recombination and, thus, fail to form chiasmata that help stabilize the homologs at the metaphase plate during meiosis I [29]. In the fruit fly, the distributive system is especially important for meiotic segregation of the fourth chromosome, which is essential for normal development but is mostly heterochromatic and does not undergo recombination [30]. Additionally, the X chromosome often relies on the distributive system for meiotic segregation since its single, large arm occasionally fails to establish a crossover between homologs, an error that rarely occurs for chromosomes 2 and 3, which each have two large arms. Considering our findings, it is possible that *N. vitripennis* females do not have a distributive system that would otherwise ensure the nuclear retention and segregation of PSR. In support of this idea, loss-of-function mutations in the *D. melanogaster* gene *nod*, a kinesin-like protein that is essential for the distributive system, results in a similar loss of the fourth chromosome from meiotic nuclei [31]. In principle, *N. vitripennis* may be able to do without a distributive system because there are no primarily heterochromatic chromosomes in the wasp’s essential genome.

Another possibility is that PSR loss may result from the observed absence of a spindle in mature *N. vitripennis* eggs. In these cells, the nuclear material appears condensed, indicating entry into meiosis I, but no microtubules are visible at this time. This pattern stands in contrast to the mature eggs of *D. melanogaster*, which are arrested during metaphase of meiosis I and have a well-formed spindle. The spindle-less *N. vitripennis* eggs are reminiscent of primordial oocytes in mammals, which are formed before birth and held in a prophase I state until puberty, when meiosis resumes to meiosis II upon ovulation and finishes upon fertilization [32]. In *N. vitripennis*, well-formed spindles were observed in freshly laid eggs, indicating that activation, likely triggered by egg laying, causes spindle formation and progression through the remainder of meiosis. Without the presence of spindle fibers in mature, unactivated wasp eggs, PSR may lack the ability to remain associated with the nuclear mass. The essential chromosomes may be stabilized within the nuclear mass at this time due to homolog pairing and crossing over.

The second aspect of B inheritance examined here is whether PSR’s copy number can increase above one. The answer appears to be no. From crosses between PSR+ parents, the proportion of F1 progeny carrying two PSR copies at the early embryonic stage was remarkably close to the expected proportion. However, there were no adult F1 progeny from these same parents that contain more than one copy. Two possible explanations may explain this pattern. First, the individuals that contain two PSR copies as embryos may die during development. A previous study showed that PSR expresses ∼70 genes in the testis, and one of these genes, *haploidizer*, plays an essential yet mysterious role in PGE [21]. It is possible that *N. vitripennis* is dosage sensitive to the expression of *haploidizer* or perhaps other PSR-expressed genes, which may reach toxic levels in somatic cells when expressed at twice their normal levels from two B copies. Such dosage sensitivity is reminiscent of sex-linked genes in other organisms; these genes must undergo dosage-compensation to mitigate this effect [33,34]. In this scenario, wasp embryos with two PSR copies would be viable because they have not yet undergone zygotic genome activation, which occurs in most organisms during or just after the mitotic cleavage divisions [35]. An alternative possibility is that precisely one of the two PSR copies is lost or dispelled in each nucleus sometime between early embryogenesis and formation of the adult germ line. Given the unlikelihood of this possibility and the precedence of gene dosage sensitivity, the lethality model seems more plausible.

Together, these findings provide a more complete understanding of PSR transmission. The most visible element of PSR’s inheritance is its PGE activity, which converts female-destined (*i.e*., fertilized) eggs into PSR-carrying males. PGE is a potent mechanism for B drive given the high rate of fertilization in *N. vitripennis*. The meiotic drag effect revealed here is normally masked by the strong female-to-male converting effect; as a result, PSR is rarely present in females, and thus not normally transmitted by them. However, PSR+ females are expected to arise naturally from spontaneous loss-of-function mutations in *haploidizer*. The meiotic drag effect would hinder the transmission of such non-driving PSR variants, thus helping to maintain strongly driving Bs in the population. Meiotic drag would also provide fitness benefits to the wasp by reducing the frequency of doomed progeny having two PSR copies. Broadly, our work reveals unforeseen properties of PSR and the organism’s reproductive biology that help govern the successful inheritance of this extreme selfish genetic element.

## MATERIALS AND METHODS

### Perpetuation of the PSR line

The PSR chromosome was kept in the wild type wasp line, *AsymC*, which is derived from the *Labii* strain that was antibiotically cured of bacterial symbionts [36]. In each generation, PSR-carrying *AsymC* males were crossed pairwise with virgin *AsymC* females and set on *Sarcophaga bullata* hosts (Carolina Biological Supply, catalog # 173486) for 2-3 days at 25°C. After F1 adult emergence, all-male broods were kept for further propagation while female-biased broods were tossed.

### The production and genotyping of PSR-carrying females

PSR+ females were generated using a procedure that has been previously described in detail [21]. In essence, an ∼800 bp region of the full length *haploidizer* cDNA was amplified with primers containing the T7 viral promoter sequence. The resulting PCR product was column-purified using the Qiaquick PCR purification kit (Qiagen, catalog # 28104) and used as a template to perform bidirectional transcription with the Invitrogen MEGAscript RNAi kit (ThermoFisher Scientific, catalog # AM1626). Following clean-up, the dsRNA was microinjected into *AsymC* PSR+ male pupae in the yellow body and red-eyed developmental stage. Once these RNAi-treated males enclosed as adults, they were crossed pairwise with *AsymC* females. Individual broods were screened for the appearance of F1 females. To confirm that the females carry PSR, genomic DNA was extracted from a few of these individuals in each female-containing brood using the DNeasy Blood and Tissue kit (Qiagen, catalog# 69504). PCR using primers specific for *haploidizer* were used under standard amplification conditions and 50-fold diluted gDNA as a template. The sequences of these primers are: (4317-Forward) 5’-GCG ACA GCC ACC GAA TTT AC-3’ and (4317-Reverse) 5’-GAC GTG CAA AAA CCT GCA TCT-3’.

### Fitness testing of PSR+ females

The fitness of *AsymC* PSR+ females was measured by assessing four characteristics – body size, longevity, F1 egg hatch rate, and F1 adult progeny produced per female – in comparison to age-controlled, wild type *AsymC* females. All four characteristics were measured from the same batch of PSR+ or wildtype females to normalize against erroneous variation among broods. Upon eclosion, females were fed a mixture of 50% honey in water and allowed to feed on fresh *Sarcophaga bullata* pupae for 24 hours. Females were temporarily immobilized with CO_2_, and digital images were taken of each female at the same magnification using a dissecting microscope. The length from the tip of head to the tip of abdomen was measured using Adobe Photoshop. The females were then individually placed into separate vials, and each was given a fresh *S. bullata* pupa. They were allowed to oviposit into the hosts for 48 hours and switched to new hosts twice in successive 48-hour periods. Hosts from the first set were opened immediately following female removal and the number of eggs was counted. They were then incubated at 25°C for 24 hours to score the number of eggs that did and did not hatch into larvae. The second and third sets of oviposited hosts were incubated at 25°C until the F1 progeny emerged as adults. The number of F1 progeny were scored for each pupa. For longevity assessment, the same females were kept on hosts (changed to fresh hosts every 3 days) at 25°C and monitored every day until life expiration.

### Wasp embryo collection and fixation

Mature wasp eggs were collected by dissecting out whole ovaries from gravid females in 1xPBT buffer. The mature eggs were removed with ultrafine forceps from the rest of the ovary tissue and chemically preserved for exactly 30 min in a mixture of 3 mL heptane, 1.5 mL 1xPBS, and 600 uL 37% formaldehyde. The fixative was removed, and the preserved eggs were washed twice with 1xPBT to remove any traces of the fix solution. The eggs were then placed in a droplet of 1xPBT on the surface of a plastic Petri dish lid and lanced in half with a 30-gauge hypodermic needle (this step is necessary for allowing penetration of staining reagents described below). To collect young embryos, gravid female wasps were allowed to oviposit into the anterior end of host pupae for 1.5-2 hrs. Embryos were carefully collected from the hosts using an egg pick (a single side of a pair of forceps) and fixed in the same fixative solution as used for eggs. The embryos were removed from the fix solution with a pipette tip and transferred onto a small piece of Whatman paper, blotted dry, and adhered to double-sided clear tape in the bottom of a small plastic Petri dish. The eggs were hydrated with ∼1 mL of 1xPBT and rolled out of their vitelline envelopes using a 30-gauge needle. The devitellinized embryos were then transferred into a microfuge tube for staining.

### Antibody staining, fluorescence in situ hybridization, and confocal microscopy of whole mount tissues

To perform antibody staining, fixed tissues were incubated overnight in primary antibodies at 4°C under slow rotation. After three 10 min washes in 1xPBT, tissues were counterstained on a platform rocker at room temperature with Cy3-conjugated secondary antibodies for 1 hr. The tissues were then washed 3 times for 10 min in 1xPBT and mounted in Vectashield mounting medium containing 4’, 6-diamidino-2-phenylindole (DAPI) (Vector Laboratories, H-1200-10). To perform DNA fluorescence in situ hybridization (FISH) on tissues that have been antibody-stained, the tissues were post-fixed in 4% paraformaldehyde in 1xPBT for 30 min at room temperature and washed 2x in 2xSSCT buffer. The tissues were then treated with a series of 2xSSCT containing increasing amounts of formamide, followed by an overnight incubation in 1.1x hybridization buffer with diluted probe. Subsequently, tissues were washed in the same series of 2xSSCT with formamide in decreasing amounts, washed twice with 2xSSCT, and mounted on a slide. The probe used to visualize PSR was synthesized and conjugated with Cy3 by IDT DNA, Inc.; the sequence is: 5’-CAC TGA AAA CCA GAG CAG CAG TTG AGA-3’. This description of DNA FISH captures the major steps of a more detailed, previously published protocol [37]. Image collection was performed using a Leica SPE DMIRB inverted fluorescence confocal microscope. Most images represent collapsed Z series to capture cellular objects in different fields of view. Images were exported as JPEG files for figure-building using Adobe Photoshop.

### Preparation and microscopic analysis of squashed sperm cells

To determine the copy number of PSR in mature sperm cells, we dissected reproductive tracks from F1 males produced from crosses between two PSR+ parents. These tissues were fixed in a 15 uL drop of 45% acetic acid and 2.5% paraformaldehyde on a cover slip for exactly 4 minutes and then squashed between the cover slip and a microscope slide. After freezing the slide in liquid nitrogen, the cover slip was quickly flicked off at one of its corners with a razor blade, soaked in 100% ethanol for 10 min, and air dried. For DNA FISH, the PSR probe (see above) was diluted to 100 ng/uL in 1.1x hybridization buffer and placed onto the tissue spot on the slide, and a cover slip was added. The slide was then heated on a hot block for 5 min at 95°C and then incubated overnight at 30°C in a humidity chamber. Subsequently, the cover slip was removed, and the slide was washed 3 times in 2xSSCT before mounting in Vectashield. Imaging of the mature sperm cell nuclei was performed on a Leica DM4000 B epifluorescence microscope. Collected images were exported as JPEG files and processed using Adobe Photoshop 2024.

## ACKNOWLEDGEMENTS

We are grateful to Stacey Hanlon and to the late Scott Hawley for helpful comments on earlier versions of this manuscript, especially regarding implications of our findings. This work was supported by a grant from the U. S. National Science Foundation (MCB-2127460) awarded to P. M. Ferree.

---

**Figure S1. Depiction of PSR segregation during meiosis and the first embryonic mitotic division under an alternative scenario.** Half the time, PSR would be predicted to segregate to the egg nucleus and one polar body as portrayed in Figure 3A. However, another outcome is that PSR would instead segregate into two of the three polar bodies. In this latter scenario, the egg nucleus and, thus the cleavage nuclei, would not contain PSR and the individual will not carry PSR. These two scenarios – one with PSR being transmitted to progeny and the other with progeny not having PSR – are equally likely so long as PSR undergoes normal segregation.

